# Population structure of *Drosophila suzukii* and signals of multiple invasions into the continental United States

**DOI:** 10.1101/2021.03.14.435345

**Authors:** Kyle M. Lewald, Antoine Abrieux, Derek A. Wilson, Yoosook Lee, William R. Conner, Felipe Andreazza, Elizabeth H. Beers, Hannah J. Burrack, Kent M. Daane, Lauren Diepenbrock, Francis A. Drummond, Philip D. Fanning, Michael T. Gaffney, Stephen P. Hesler, Claudio Ioriatti, Rufus Isaacs, Brian A. Little, Gregory M. Loeb, Betsey Miller, Dori E. Nava, Dalila Rendon, Ashfaq A. Sial, Cherre B. da Silva, Dara G. Stockton, Steven Van Timmeren, Anna Wallingford, Vaughn M. Walton, Xingeng Wang, Bo Zhao, Frank G. Zalom, Joanna C. Chiu

## Abstract

*Drosophila suzukii*, or spotted-wing drosophila, is now an established pest in many parts of the world, causing significant damage to numerous fruit crop industries. Native to East Asia, *D. suzukii* infestations started in the United States a decade ago, occupying a wide range of climates. To better understand invasion ecology of this pest, knowledge of past migration events, population structure, and genetic diversity is needed. To improve on previous studies examining genetic structure of *D. suzukii*, we sequenced whole genomes of 237 individual flies collected across the continental U.S., as well as several representative sites in Europe, Brazil, and Asia, to identify hundreds of thousands of genetic markers for analysis. We analyzed these markers to detect population structure, to reconstruct migration events, and to estimate genetic diversity and differentiation within and among the continents. We observed strong population structure between West and East Coast populations in the U.S., but no evidence of any population structure North to South, suggesting there is no broad-scale adaptations occurring in response to the large differences in regional weather conditions. We also find evidence of repeated migration events from Asia into North America have provided increased levels of genetic diversity, which does not appear to be the case for Brazil or Europe. This large genomic dataset will spur future research into genomic adaptations underlying *D. suzukii* pest activity and development of novel control methods for this agricultural pest.

## INTRODUCTION

Over the past decade, *Drosophila suzukii* (Matsumura), also known as the spotted-wing drosophila or the Asian vinegar fly, has become an incredibly invasive pest species and a threat to soft fruit agriculture worldwide (dos Santos *et al*. 2017). Unlike the large majority of Drosophilidae (Diptera), which preferentially breed in decaying plant material, female *D. suzukii* developed their own ecological niche by evolving a serrated ovipositor, enabling them to lay eggs in fresh ripening soft-skinned fruits (Walsh *et al*. 2011; Walton *et al*. 2016). First described in Japan as an agricultural pest of cherries, *D. suzukii* was primarily distributed across East Asia until researchers found wild specimens in Hawaii in 1980 (Peng 1937; Kanzawa 1939; Kaneshiro 1983). In 2008, *D. suzukii* was detected California, and by 2009 was widespread across the Western U.S. coast (Hauser *et al*. 2009; Bolda *et al*. 2010). In the Eastern U.S., *D. suzukii* first appeared in Florida in 2009 (Steck *et al*. 2009), before again rapidly spreading across the entire East coast within a few years. Meanwhile in Europe, *D. suzukii* was first detected in Spain and Italy in 2008 and rapidly spread across Europe, appearing in France, Switzerland, Austria, Germany, and Belgium by 2012.

Subsequently *D. suzukii* arrived in South America when it was detected in Brazil in 2013 (Deprá *et al*. 2014), Argentina in 2014 (Cichon *et al*. 2015), and Chile in 2015 (Medina-Muñoz *et al*. 2015). Its rapid spread across continents suggests that human transportation is likely a major factor, as eggs laid in fresh fruit are difficult to detect before shipment. Once established in a new continent, *D. suzukii* rapidly disperse to neighboring regions, aided by its ability to adapt to a wide range of climates through phenotypic plasticity (Shearer *et al*. 2016). In the Western U.S. Coast alone, estimated economic losses were as high as 511 million dollars per year, assuming a 20% average yield loss (Bolda *et al*. 2010). Thus, there is much interest in understanding the patterns of migration and origin of these invasive populations, as these data can be used to inform shipping and quarantine policies and to identify routes of entry.

Previous research on the population genomics of *D. suzukii* was performed using a relatively small number of molecular markers. Adrion et al. (2014) used six x-linked gene fragments from flies collected across the world, and detected signals of differentiation between European, Asian, and U.S. populations. However, they found no evidence of differentiation within the 12 U.S. populations sampled, possibly due to the limited power provided from a small number of markers. A follow-up study using 25 microsatellite loci of samples collected between 2013-2015 greatly improved estimations of migration patterns worldwide; the authors found evidence for multiple invasion events from Asia into Europe and the U.S. as well as an East-West differentiation in the 7 populations sampled in the continental U.S. (Fraimout *et al*. 2017). However, using microsatellites alone may miss more subtle signals of population structure compared to genome-wide datasets, as increasing the number of independent loci genotyped increases accuracy of population parameter estimates, even when the number of biological samples is low (Trask *et al*. 2011; Willing *et al*. 2012; Rašić *et al*. 2014). With the advent of affordable whole-genome sequencing, it has become feasible to sequence hundreds of individuals to study population genomics, enabling improved inference of population structure using hundreds of thousands to millions of SNP markers (Soria-Carrasco *et al*. 2014; Wu *et al*. 2019; Lee *et al*. 2019).

In this study, we leverage the power of whole-genome sequencing to individually sequence hundreds of *D. suzukii* samples to determine whether U.S. populations are stratified along a North-South cline corresponding to varying winter climates, as well as to detect whether migration is freely occuring between the East and West Coasts. In addition, we include several representative populations from Asia, Europe, and Brazil to determine frequency and source of international migrations and compare genetic diversity between invasive and native populations. We expect these analyses and the large sequencing dataset will be of value in developing policies and furthering research into mitigating the harmful effects of *D. suzukii* worldwide.

## MATERIALS AND METHODS

### Sample collection and genomic DNA extraction

We received either flash-frozen or ethanol-preserved samples of *D. suzukii* for genomic analysis. Japanese samples were a lab strain from the Kyoto Japanese Stock Center; Hawaiian samples were wild-caught in 2009 and kept as a lab colony until DNA extraction in 2017; all other samples were field-collected. Ethanol-preserved samples were re-hydrated in 100uL water prior to DNA extraction. Flies were individually disrupted using a 3mm diameter steel bead in a TissueLyser (Qiagen, Germantown, MD) for 30 seconds at 30Hz in 100uL of 2mg/mL Proteinase K in PK buffer (MagMAX™, Thermofisher Scientific, Pleasanton, CA) before being spun down in a centrifuge for 1 minute at 10,000rpm and incubated for 2 hours at 56°C. 100uL of MagMAX DNA lysis buffer was added to each sample, followed by a 10 minute incubation, before proceeding to DNA purification using a BioSprint DNA Blood Kit on a BioSprint 96 Workstation (Qiagen), using protocol “BS96 DNA Tissue” as per manufacturer’s instructions. Supplemental Table 1 contains all sample names, collection locations, and time of collection.

### Library preparation and sequencing

Illumina sequencing libraries were prepared using either the Kappa HyperPlus Kit (Roche, South San Francisco, CA) (lanes 2-4) or Qiaseq FX DNA Library Kit (Qiagen) (lanes 5-8) using 50 ng of input DNA. We followed the manufacturer’s instructions for both library preparation kits with few exceptions. With the Kappa HyperPlus Kit, we fragmented DNA at 30°C for 20 minutes and increased adapter incubation time to 1 hour. We also added a 0.6X and 0.7X size selection with AmPure XP beads (Beckman Coulter Life Sciences, Indianapolis, IN) following 5 cycles of PCR amplification with an Eppendorf Master Cycler Pro (ThermoFisher Scientific). With the Qiagen FX kit, we fragmented DNA at 30°C for 15 minutes, and amplified with 7 cycles of PCR. In both cases, DNA library concentration and fragment size were quantified on a Qubit (ThermoFisher Scientific) and a Bioanalyzer High-Sensitivity DNA chip (Agilent, Santa Clara, CA). Paired-end 150 base-pair sequencing was performed by Novogene, Inc. (Sacramento, CA) on the Illumina HiSeq 4000 platform.

### Alignment to D. suzukii reference genome

Raw Illumina reads were inspected for quality using fastqc version 0.11.5 (Babraham Institute, Cambridge, UK), and trimmed for low quality bases and adapter sequences using Trimmomatic version 0.35 (Bolger *et al*. 2014), using the following parameters: Leading qscore threshold = 10, trail score threshold = 10, minimum read length = 36, and illuminaclip=2:30:10. Reads were then aligned to the *D. suzukii* reference genome (genbank accession GCA_000472105.1) (Chiu *et al*. 2013) using bwa-mem version 0.7.9a (Li 2013), sorted by samtools-sort verson 1.3.1(Wellcome Trust Sanger Institute, London, UK), de-duplicated with picardtools-MarkDuplicates version 2.7.1 (Broad Institute, Cambridge, MA), and indexed with samtools-index version 1.3.1. Variants were called using freebayes version 1.1.0 (Garrison and Marth 2012) with default parameters and filtered to only include variants with QUAL>30.

### Analysis of D. suzukii, D. pulchrella, and D. subpulchrella COX2

The *D. suzukii* mitogenome sequence and ten *D. pulchrella* COX2 sequences were downloaded from NCBI. *Drosophila subpulchrella* mitogenome was identified by running BLAST with the *D. suzukii* mitogenome against the *D. subpulchrella* genome assembly (GCA_014743375.2), and annotated using MITOS2 (Bernt *et al*. 2013). COX2 sequences from all *D. suzukii* samples were identified by aligning raw reads to the *D. suzukii* mitogenome, filtering out any read pairs where one of the reads was unmapped (samtools view –f 2 –F 4). Variants were called with freebayes version 1.1.0 in haploid mode, and fasta sequences were extracted with bcftools-consensus version 1.10.2 (Wellcome Trust Sanger Institute).

All COX2 sequences were aligned with the ClustalOmega web portal (Madeira *et al*. 2019), and haplotypes were called using DNASP version 6.12/03 (Rozas *et al*. 2017). The evolutionary history was inferred by using the Maximum Likelihood method and General Time Reversible model (Nei and Kumar 2000) in MEGA version 10.1.8 (Kumar *et al*. 2018). Initial trees for the heuristic search were obtained automatically by applying Neighbor-Joining and BioNJ algorithms to a matrix of pairwise distances estimated using the Maximum Composite Likelihood (MCL) approach, and then selecting the topology with superior log likelihood value. A discrete Gamma distribution was used to model evolutionary rate differences among sites (5 categories (+*G*, parameter = 0.1066)). The tree is drawn to scale, with branch lengths measured in the number of substitutions per site. This analysis involved 21 nucleotide sequences from a total of 595 positions in the final alignment.

### Genotype clustering and discriminant analysis

Variants located on scaffolds shorter than 100kb were excluded to avoid potentially misassembled scaffolds. As X-linked and autosomal markers are expected to have different genetic diversity levels and effective population sizes, and samples were not sexed, we decided to focus analysis on autosomal markers. X-linked scaffolds were identified and removed by aligning our reference genome using nucmer version 4.0 (Marçais *et al*. 2018) against known X-linked scaffolds from an annotated *D. suzukii* PacBio genome assembly (Paris *et al*. 2020).

To decide on filtering cutoffs for downstream Discriminant Analysis of Principal Components (DAPC) and Principal Component Analysis (PCA), VCFtools version 0.1.14 was used on a 10% random subset of the VCF file to inspect distribution of minor allele frequency, fraction of missing genotypes per site, read depth, and fraction of genotypes missing per sample (Supp. Figure 1). The fraction of missing genotypes within each sampling location was also inspected to exclude loci that were missing data within a sample location. Based on visual inspection of the plotted data, the following parameters were used to filter loci: only include biallelic SNPs, minimum minor allele count = 5, maximum read depth per genotype = 100, maximum mean depth per site = 40, minimum read depth per genotype = 3, minimum mean depth per site = 5, maximum missing genotypes per site across all samples = 10%, and maximum missing genotypes per site within each sampling location = 40%. Samples DD1-4 and WA1-5 were excluded from the analysis due to their high amount of missing genotype data. Loci were then pruned for linkage disequilibrium using plink/1.9 (Chang *et al*. 2015) with a window size of 50 SNPs, stepping 5 SNPs at a time, using a variance inflation factor (VIF) threshold = 2, corresponding to a multiple correlation coefficient less than 0.5. This left 106,766 loci for analysis.

The R package “adegenet” version 2.1.3 (Jombart and Ahmed 2011) was used to cluster samples, calculate principal components, and perform discriminant analysis. Sample clustering was conducted for k=1 to k=20, and a Bayesian Information Criterion score was calculated to determine which values of k best fit the data. A-score optimization was run to determine the optimal number of PCs to include in DAPC for each value of k. Posterior probabilities of assignment for multiple values of k were plotted using CLUMPAK (Kopelman *et al*. 2015).

### Treemix migration trees and F3/F4 statistics

Filtering of the VCF was performed with the same cutoffs as for DAPC analysis, except for the fraction of missing genotypes within each sampling site. Instead, samples were grouped by region as follows: West Coast plus Alma Research Farm, Bacon county, GA (AR); Hawaii; Brazil; East Coast (without AR); Europe; Sancheong, South Korea (SN); Namwon, South Korea plus Kunming, China (KM_NW); and Japan. Loci were filtered out if missing greater than 15% data within these groups. In addition, sample SN1 was excluded from the dataset as based on DAPC this sample was an outlier in the SN population, and could have represented a mislabeled sample. This left 265,699 SNPs for tree estimation after filtering. Treemix version 1.13 (Pickrell and Pritchard 2012) was used to generate maximum likelihood trees. Between 0 and 5 migrations were allowed, with 100 bootstraps calculated using a resampling block size of 500 SNPs. The bootstrap run with maximum likelihood for each migration tested was used to for plotting and viewing of migration edges. Treemix was also used to calculate F3 and F4 statistics for all combinations of regions, using jackknife estimation with a block size of 500 SNPs to estimate standard error and Z-scores. The F3 statistic tests if population A’s allele frequencies are a result of mixture of allele frequencies from populations B and C. A significantly negative value of F3(A;B,C) supports admixture of B and C into A. The F3 statistic measures correlations in allele frequencies between populations A and B versus populations C and D. F4(A,B;C,D) is expected to be zero under no admixture. A significantly positive value suggests gene flow between A and C or B and D, while a significantly negative value suggests gene flow between B and C or A and D.

### Estimation of F_ST_, nucleotide diversity, and Tajima’s D

Pairwise weighted F_ST_ values were calculated using ANGSD version 0.933 (Korneliussen *et al*. 2014) from BAM files for scaffold1 of the assembly (approximately 22.9 Mb). To do this, the site allele frequency for each group was estimated using the following parameters: -doSaf 1 -GL 1 -baq 1 -C 50 -minQ 20 -minmapQ 30. Next, the 2D site frequency spectrum for all pairs of groups was estimated using ANGSD/realSFS. Finally, F_ST_ was estimated at each locus using “ANGSD/fst index”, and a global F_ST_ for each pair was calculated using “ANGSD/fst stats”. Nucleotide diversity statistics were calculated using ANGSD from BAM files across the entire genome. Site allele frequency and 2d-SFS were calculated the same way as for F_ST_. Nucleotide diversity and Tajima’s D were estimated per site using ANGSD/saf2theta, and then binned in 20kb windows across the genome using ANGSD/thetaStat do_stat - win 20000.

## RESULTS

### Population structure exists between continents as well as within the U.***S. and Asian populations***

To identify whether population structure exists in *D. suzukii* living in recently invaded locations, we sampled and sequenced wild caught individual *D. suzukii* flies collected from the continental United States, Brazil, Ireland, Italy, South Korea, and China, as well as a laboratory strain from Hawaii and Japan (Figure 1, Supp. Table 1). After aligning sequences to the reference genome and calling single nucleotide variants, we performed PCA and DAPC. The first four principal components (PCs) explaining allelic variation were informative to identify population structure (Figure 2A). The first PC, accounting for 6.2% of variance in the data, clearly separates an Asian cluster consisting of KM (China) and NW (South Korea) from the remaining samples, while the second component broadly separates the remaining samples by geographic origin. Surprisingly, nearly all Sancheong, South Korea samples (SN) cluster more closely with invasive European and New World samples than to NW, despite being collected from a nearby location on nearly the same date. In addition, the Japanese lab strains appear to be distinct from other samples, likely due to strong genetic drift due to its long duration in captivity.

**Figure 1:**
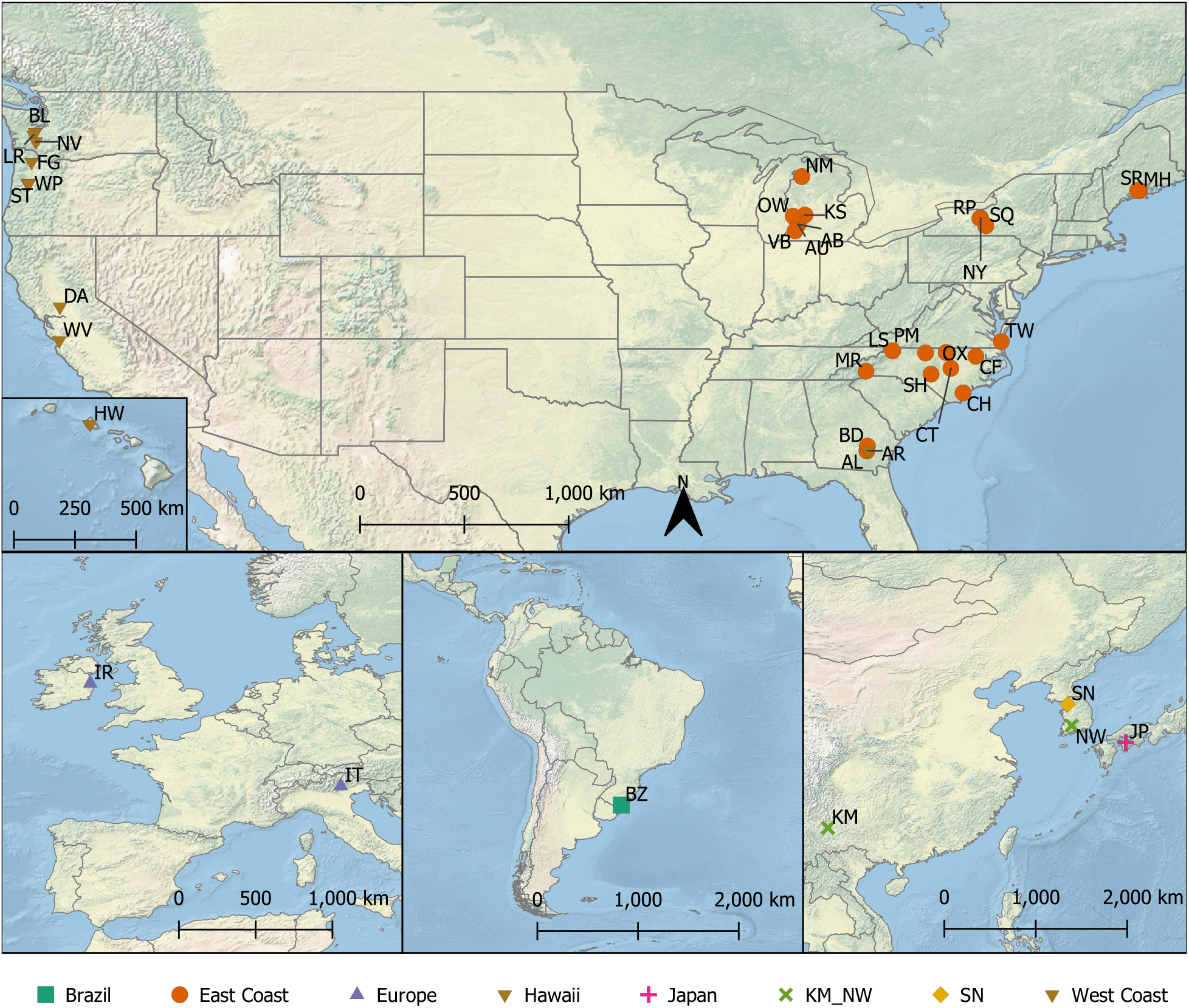
Sampling sites of *Drosophila suzukii* populations. Sampling sites from the United States, Europe, Brazil, and Asia. Colors and symbols depict population clusters identified by DAPC and used in downstream analyses. Between 5-10 flies per site were collected for whole-genome sequencing. Refer to Supp. Table 1 for details of collection sites. AB=USA:Allegan, MI. AL=USA:Wade Lane, Alma, Bacon County, GA. AR=USA:Alma Research Farm-Bacon County, GA. AU=USA:Allegan, MI. BD=USA:Bazley Dole Farm, Appling County, GA. BL=USA:Olympia, WA. BZ=Brazil:Pelotas, Rio Grande Do Sul. CF=USA:Cherry Farm, NC. CH=USA:Castle Hayne, NC. CT=USA:Clayton, NC. DA=USA:Davis, CA. DD=China:Dandong. FG=USA:Banks, OR. HW=USA:Oahu, HI. IR=Ireland:North Dublin. IT=Italy:Trento, Pergine Valsugana. JP=Japan:Ehime. KM=China:Kunming, Yunnan. KS=USA:Kent, MI. LR=USA:Little Rock, WA. LS=USA:Laurel Springs, NC. MH=USA:Marrs Hill Rd-Union, ME. MR=USA:Mills River, NC. NM=USA:Traverse, MI. NV=USA:Napavine, WA. NW=South Korea:Namwon. NY=USA:Geneva, New York. OW=USA:Ottawa, MI. OX=USA:Oxford, NC. PM=USA:Piedmont, NC. RP=USA:Ontario County, NY. SH=USA:Sandhills, NC. SN=South Korea:Sancheong. SQ=USA:Schuyler County, NY. SR=USA:Sidelinger Road, Union, ME. ST=USA:53rd Street-Corvallis, OR. TW=USA:Virginia Beach, VA. VB=USA:Van Buren, MI. WA=USA:Watsonville, CA. WP=USA:Willamette Park-Corvallis, OR. WV=USA:Watsonville, CA.

**Figure 2:**
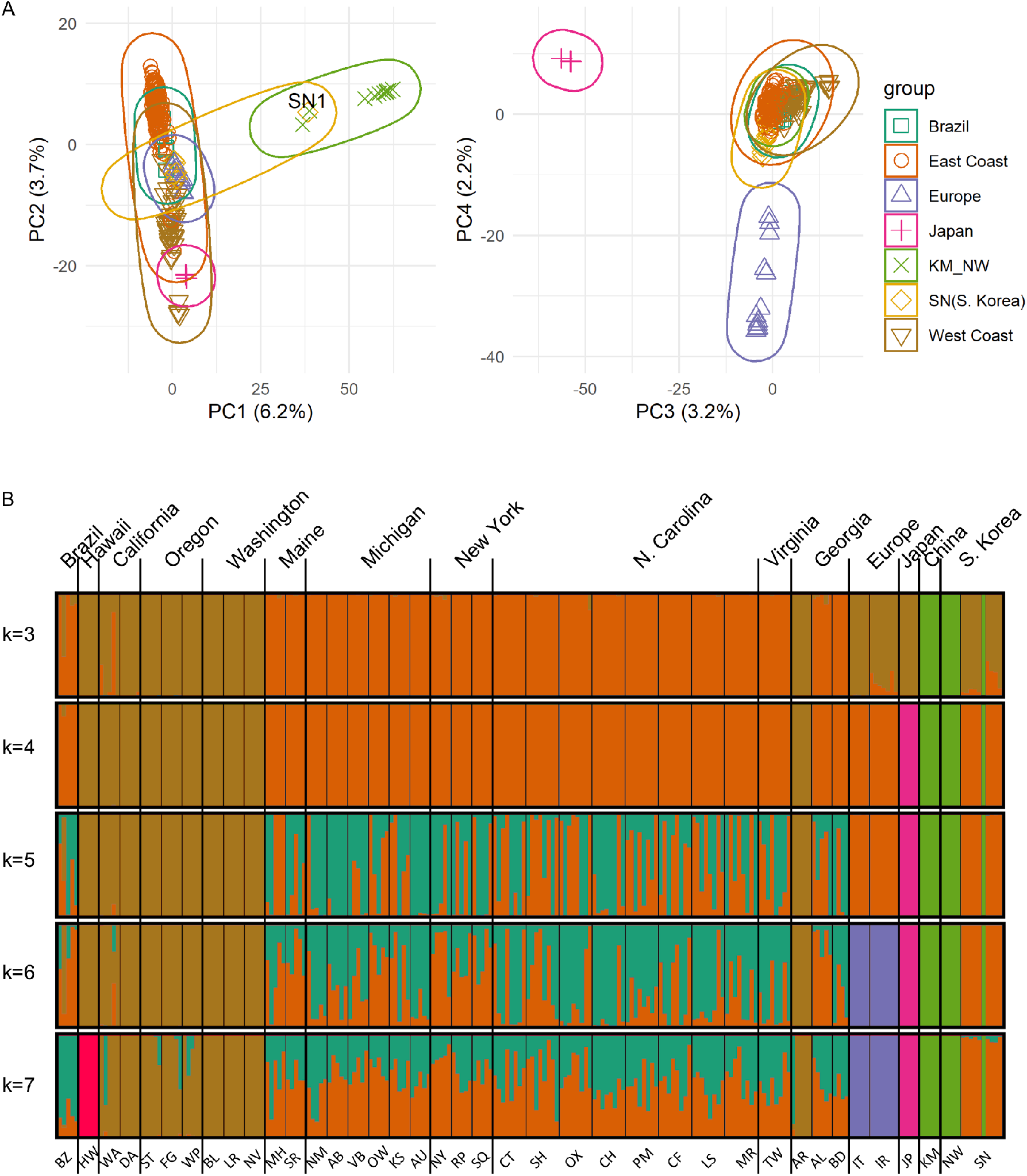
Population Structure of *Drosophila suzukii* populations. **(A)** First 4 principal components plotted of the sampled regions based on 106,766 SNPs. Note the SN1 outlier in PC1 vs PC2 plot. Percent variation of the data captured by each component indicated in axis labels. **(B)** Posterior probability of cluster identity using Discriminant Analysis of Principal Components calculated from 106,766 SNPs, using between 3 to 7 clusters. Top labels indicate broader region while bottom labels correspond to specific sampling sites (see Supp. Table 1).

To evaluate more fine-scale differences between individuals and populations, we used discriminant analysis of principal components to estimate posterior probabilities of assignment to a given number of clusters(k) between 3 and 7 (Figure 2B). At each level of k, we gained some insight into differences between individuals. At all levels of k, we observed that KM and NW were distinct from all other populations, matching our observations in the first principal component. To ensure that the strong differentiation of KM and NW from the other samples was not due to misidentification of species, we aligned the COX2 gene sequence from morphologically similar *D. suzukii, D. pulchrella*, and *D. subpulchrella*, along with all our sequenced samples, to infer a phylogenetic tree (Supp. Figure 2). Both *D. pulchrella* and *D. subpulchrella* have long branches on the tree relative to all our sequenced samples and the *D. suzukii* reference mitogenome, confirming all our samples are *D. suzukii*.

We also observed that the Western and Eastern U.S. samples are distinct and relatively non-divisible, suggesting they are separate populations with gene flow occurring freely within these regions. The one exception to this is the flies sampled from Alma Research Farm, Bacon County, Georgia (population AR). They appear strongly “West Coast” in ancestry, which could represent a recent or isolated migration to the area. Based on this, we grouped AR with the West Coast population for the rest of the analyses. We also observed that Hawaiian samples clustered tightly with West Coast populations for most values of k. As Hawaii was invaded by *D. suzukii* much earlier than the West Coast, this supports the theory that West Coast populations owe some of their ancestry to Hawaiian introgression. However, at k=7 we see Hawaiian samples form their own distinct cluster, likely a result of genetic drift or its time in captivity prior to sequencing. As a result, we grouped Hawaiian samples with West Coast samples for the remainder of the analyses. Of the five individuals sampled from Pelotas, Rio Grande do Sul, Brazil, most appear “East Coast” in origin, with one individual potentially carrying some “West Coast” ancestry, suggesting Brazil is admixed from both U.S. coastlines. At higher values of k, the populations from Europe appear isolated as an individual cluster as well, validating their distinctness from other invasive populations we see from principal component plotting.

Interestingly, at different values of k, SN clusters closer with “New World” populations than other Asian populations or Europe. At k=3, SN and the West Coast form a single group, while at higher values of k, SN becomes its own cluster (blue) and appears to represent a large portion of the East Coast’s ancestry, in addition to a potentially unsampled “ghost” population (maroon). We did find one potential outlier sample, SN1, which appears to be consistently grouped with KM_NW at all values of k, as well as in all PCA plots. Based on concerns that this could be due to sample labeling error, we excluded this sample from the remaining analyses.

Finally, to quantify the amount of differentiation present between regions, we estimated Fst values between regions using the largest scaffold in the reference genome (Figure 3C). Fst values for Japan and KM_NW versus all other populations were high (>0.50), while Fst between the U.S., Europe, Brazil, and SN ranged between 0.07 to 0.17. In agreement with the PC and DAPC data, Fst values of SN are lowest between the West Coast (0.09) and East Coast (0.10), supporting the idea than SN is closely related to U.S. invasive populations.

**Figure 3:**
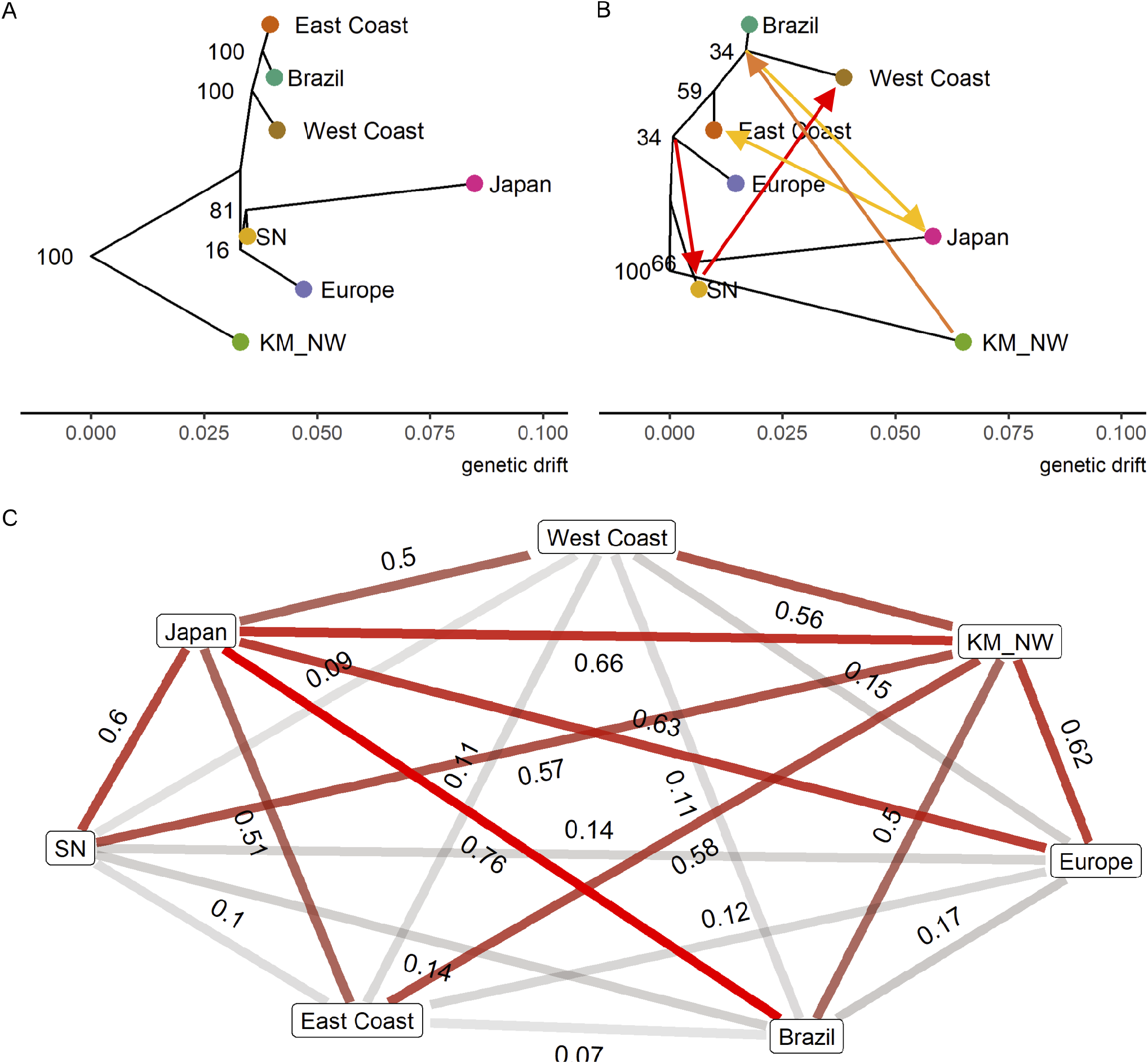
Differentiation and relationships between sampled regions. **(A)** Maximum likelihood tree based on allele frequencies without any migrations. Nodes labeled with bootstrap confidence percentages obtained from 100 replicates. **(B)** Maximum likelihood tree allowing 5 migration events. Migrations are colored by strength from red (strong) to yellow (weak), with the following admixture proportions: 0.458 SN to West; 0.455 Europe/East/West/Brazil to SN; 0.337 KM_NW to West/Brazil; 0.139 Japan to East; 0.048 Brazil to Japan. **(C)** Pairwise weighted Fst calculated from scaffold 1 of the reference genome between populations. Darker red lines indicate more differentiation between populations.

### Maximum likelihood tree analysis shows evidence of multiple invasion events from Asia into North America

While the DAPC identified clusters *de novo* and suggested potential admixture, it is unable to provide more detailed understanding of population history or migration events. In order to estimate the population history of these invasive populations, we used Treemix to generate a population tree with migration events based on allele frequency differences between the populations identified from DAPC. To test for significance of any migration events, we also calculated F3 and F4 statistics (Supp. Table 2 and Supp. Table 3). Using a SNP dataset pruned of linked sites, we tested between 0 to 5 migration edges, bootstrapping each analysis 100 times and plotting the tree with maximum likelihood (Figure 3A-B, Supp. Figure 3). We used KM_NW to root the tree, due to its high degree of differentiation from other populations and because Asia represents the ancestral home of *D. suzukii* (Peng 1937; Kanzawa 1939). When no migrations are allowed, the New World samples form a single clade, while Europe forms a clade with SN and Japan (max likelihood = 38.4353). Residuals between the treemix model and the data range between +/- 4 standard errors (Supp. Figure 4). However, upon allowing up to 5 potential migrations to flow between branches of the tree, we find several strong migration events that improve model fit (max likelihood = 228.486, residuals +/- 0.3). The strongest signal, with an admixture of 45.8%, suggests a migration from SN to the West Coast. To test this formally, we calculated F4 statistics with Brazil and the West Coast as one sister group, and either KM_NW, SN or Europe, SN as the second sister group, and found significantly positive values in both cases (Z-score 11.31 and 11.54, respectively; Supp. Table 3), supporting a Brazil/West Coast admixture event. We also see a strong migration from KM_NW to the West Coast/Brazil node, and an F4 test of the form (Brazil, East Coast; Japan, KM_NW) is significantly negative (z-score -6.534), supporting either a Brazil/KM_NW or East Coast/Japan admixture. Japan/East Coast admixture is unlikely as the Japanese lab population was obtained in 2002, years before *D. suzukii* appeared in the continental U.S. As such, these migrations likely represent multiple invasion events from Asia into the New World.

### U.S. populations exhibit relatively high diversity and little evidence of bottleneck, unlike European and Brazilian populations

To determine if invasive populations have experienced loss in genetic diversity, we used the software ANGSD to estimate average pairwise nucleotide diversity and Tajima’s D in 20 kb increments across the genome for each population. Invasive populations can sometimes exhibit reduced levels of diversity early on in their history due to a founder effect (Nei *et al*. 1975), while ancestral populations tend to have the greatest amount of diversity due to their existence in the region for many generations. As the presumed ancestral population, KM-NW exhibits the largest amount of nucleotide diversity, followed by the U.S. and SN samples with intermediate levels of diversity, followed by Brazil and Europe with relatively low levels (Figure 4A). The distribution of Tajima’s D supports a lack of bottlenecking across populations, with the exception of Europe exhibiting a relatively high average value of Tajima’s D across the genome (Figure 4B). A bottleneck of intermediate strength can cause this due to the loss of many rare alleles during the population shrink followed by rapid population growth, allowing any previously rare alleles to grow to an intermediate frequency, resulting in a negative Tajima’s D immediately following the end of the bottleneck (Tajima 1989). The lack of migration edges from the Treemix analysis, combined with low diversity levels and high Tajima’s D, suggest *D. suzukii* populations from Europe were the result of a single migration event.

**Figure 4:**
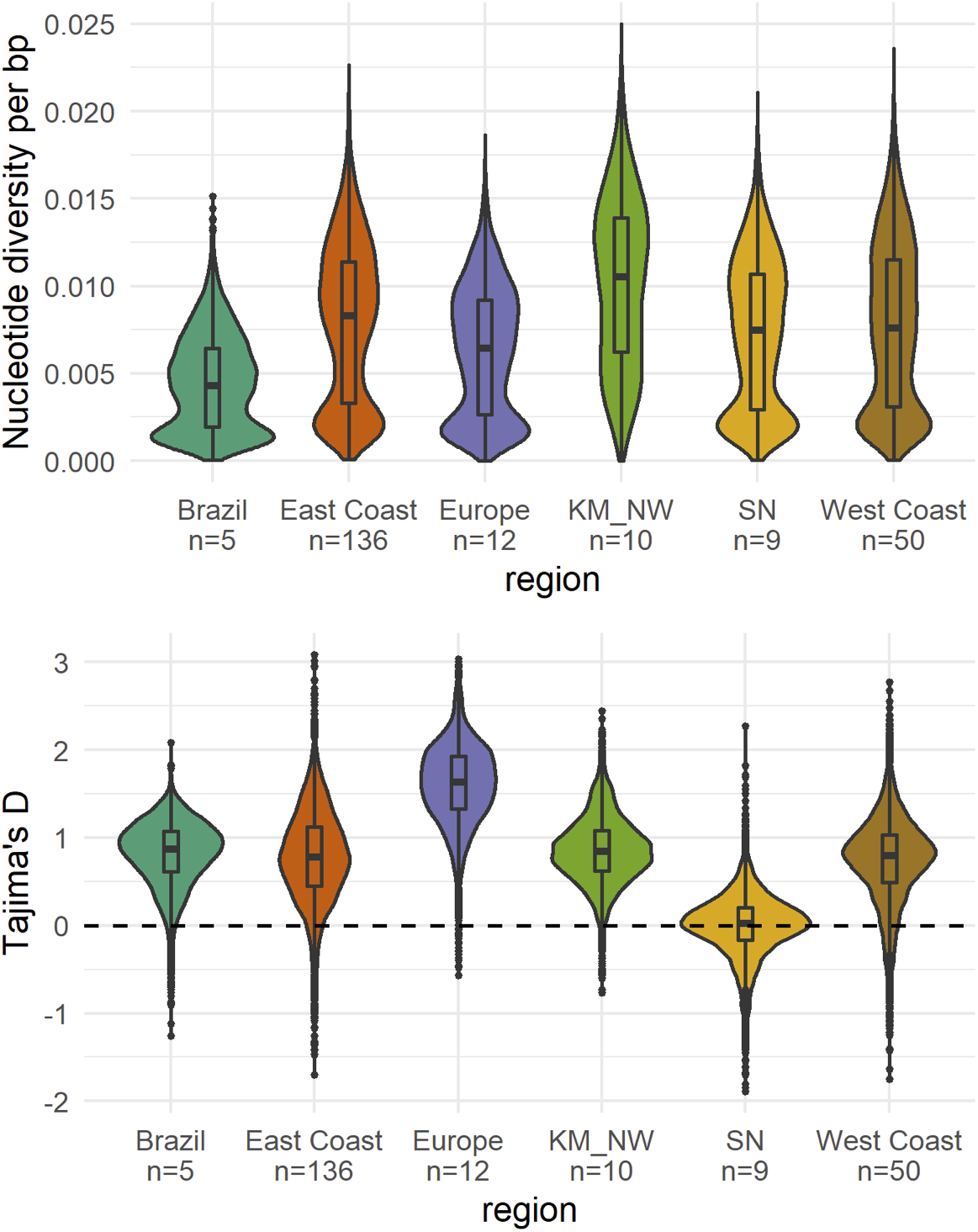
Genetic Diversity of sampled *D. suzukii* populations. **(A)** Pairwise nucleotide diversity distribution, calculated in 20 kb intervals across the *D. suzukii* genome. Boxplots depict median, 1^st^ and 3^rd^ quartiles. Number of diploid samples indicated along the x-axis. **(B)** Distribution of Tajima’s D, calculated using a sliding window of 20 kb with 5 kb steps. Boxplot depicts median, 1^st^ and 3^rd^ quartiles. Dotted line indicates expected value of 0 for a neutrally evolving stable population.

## DISCUSSION

Based on population allele frequencies, we have shown that *D. suzukii* exhibit population structure based on region and invasion history. In the New World populations, we find that Eastern and Western U.S. samples appear to be distinct populations. While this could be the result of a continuous population variation from East through Central to the West coast, it is more likely the case that the two populations experience little gene flow due to strong geographic barriers such as the Sierra Nevada or Rocky Mountain ranges, and the fact that key target fruit crops such as cherries, raspberries, blueberries, and strawberries are primarily grown in states that we sampled (“Noncitrus Fruits and Nuts 2019 Summary” 2020). Any genetic exchange between these regions would likely be the result of human activity, such as could be the case with samples collected from Alma Research Farm, Bacon County, Georgia (AR), with all samples clustering with the West Coast populations.As other nearby collections (AL, BD) failed to share this signal, the Alma research population likely represents a recent and isolated migration event. The fact that we see little evidence of regular gene admixture within the U.S. is somewhat surprising as the country’s supply of fresh blueberries, cherries and caneberries are concentrated in a few states (Pacific Northwest, Michigan, Maine) (“Noncitrus Fruits and Nuts 2019 Summary” 2020). However, recent changes to cultural management such as more frequent harvesting and post-harvest chilling may be responsible for disrupting the *D. suzukii* lifecycle and limiting cross-country transport (Schöneberg *et al*. 2021).

Several studies have reported a low probability of *D. suzukii* surviving when exposed to freezing temperatures, based on cold survival assays of wild-caught specimens (Dalton *et al*. 2011; Stephens *et al*. 2015), suggesting that flies collected in regions such as Washington, Michigan, Maine, and New York could be annual migrants to the area from nearby warmer locations. Based on our cluster analysis, there appears to be no broad level differentiation between North-South clines in either West or East Coast populations, supporting the theory that flies are regularly migrating into colder climates after the harsh winters have ended. Alternatively, flies could be tolerating winters by surviving inside human structures (Stockton *et al*. 2019), or by having evolved resistance to freezing temperatures (Stockton *et al*. 2020). Studies using *D. suzukii* collected from different locations have reported different levels of rapid cold-hardening response, suggesting there could be regional selection present (Jakobs *et al*. 2015; Everman *et al*. 2018; Stockton *et al*. 2020). In addition, the majority of the samples in the U.S. were collected in the summer; if populations in Northern regions undergo seasonal fluctuations in allele frequencies, such as has been demonstrated in wild *D. melanogaster* populations collected in Pennsylvania (Bergland *et al*. 2014), we may be overlooking structural differences that occur during the winter. Likely, some combination of these factors is responsible for the success of *D. suzukii* in these regions, and further studies will be needed to identify the causes. North-South clines in traits such as diapause and circadian rhythms have been previously identified in drosophilids and could be at play here as well (Schmidt *et al*. 2005; Tyukmaeva *et al*. 2011). Further analyses with a recently published and well annotated genome (Paris *et al*. 2020) could improve detection of genes under selection in this dataset, using methods such as those recently used to detect SNPs correlated with invasive success (Olazcuaga *et al*. 2020).

Despite being recently established invasive populations, both East and West Coast clusters exhibit nucleotide diversity levels only moderately below that of the presumed native Asian population. Typically, recent invasion events are characterized by reduced diversity due to bottleneck effects (Dlugosch and Parker 2008). The fact that U.S. samples appear quite diverse suggests either a large population initially invaded both regions, or that recurrent migration from Asia is providing genetic diversity. This increased diversity may affect the success of control attempts, as there may be higher potential for resistance development and local adaptation. Treemix’s inference of admixture events from SN, Korea into the West Coast and from KM_NW into the West Coast/Brazil cluster support the idea that Western U.S. populations have been supplemented several times post-invasion. This has also been previously suggested based on microsatellite data (Fraimout *et al*. 2017). Furthermore, values of Tajima’s D for both U.S. populations are not substantially different from KM_NW values, suggesting little to no bottleneck effects.

This is in sharp contrast to the European populations sampled, which have larger reductions in nucleotide diversity than New World populations, as observed previously (Adrion *et al*. 2014). Europe’s broadly high values of Tajima’s D also correspond to an intermediate bottleneck effect. Estimates of bottleneck severity in different worldwide populations by Fraimout et al. (2017) also corroborate this finding. Similarly, the Brazilian population displays highly reduced nucleotide diversity relative to the West Coast, suggesting that the KM_NW migration to the West Coast/Brazil node happened after *D. suzukii* were introduced to Brazil. As Europe and Brazil were not as heavily sampled as the U.S. in this study, there may be unaccounted population structure at play. A study using the mitochondrial COI sequence of *D. suzukii* from 12 Brazilian sample sites found only 5 haplotypes but did observe evidence of population structure by geography, suggesting a single invasion event with subsequent expansion (Ferronato *et al*. 2019). Alternatively, these low diversity levels, combined with the fact that no admixture events directly to Europe or Brazil were detected in Treemix, could mean that European and Brazilian populations will have limited ability to adapt, which can be beneficial for control strategies. Successive invasion/migration events can provide relief from any initial bottlenecks by providing increased genetic diversity, and has been observed to occur in multiple animal studies (Johnson and Starks 2004; Kolbe *et al*. 2004), which should lead to increased ability to adapt and evolve to new climates. Measures to reduce impacts of invasive species are often hindered by repeated introgressions or migrations into the invaded zone (Garnas *et al*. 2016). As invasive species transport is strongly associated with international trade of live plants and plant products (Chapman *et al*. 2017), it will be important to enforce that fruits from overseas regions are free of live *D. suzukii* as required by the U.S. Department of Agriculture, even though this species is already established in the U.S.

We anticipate that the hundreds of sequenced genomes provided here will prove useful in many fields of biology beyond the scope of this study. Knowledge of genetic variation and alternate alleles present within a species can be informative for the design of probes and micro RNAs (miRNAs), such as for the purpose of creating gene drives to control invasive species. Gene drive mechanisms to eliminate *D. suzukii* have been experimentally tested on multiple lines to ensure the miRNAs are broadly effective (Buchman *et al*. 2018), but a large dataset of wild population sequencing allow researchers to more confidently select target sites that are non-variable and thus non-resistant to Cas9 targeting (Schmidt *et al*. 2020). Drury *et al*., (2017) demonstrated that minor natural polymorphisms in target sites reduce gene drive effectiveness in flour beetles, and tools have been developed to help researchers design gRNAs accounting for population variation (Chen *et al*. 2020).Similarly, with the recent development of a CRISPR-Cas9 editing and RNAi knockdown protocols for *D. suzukii* (Murphy *et al*. 2016; Li and Scott 2016; Taning *et al*. 2016; Ahmed *et al*. 2020), prior knowledge of allelic variation will allow researchers to design targeting oligonucleotides more precisely to avoid loci with variability. Most recently, our dataset has been used to study sensory receptor evolution in *D. suzukii*, giving insights into its evolution toward becoming a major agricultural pest (Durkin *et al*. 2021). Other future uses of this trove of genomic data could involve insecticide resistance studies or the development of diagnostic assays for rapid detection in the field.

## DATA AND REAGENTS AVAILABILITY

Raw Illumina reads have been deposited to the NCBI Sequence Read Archive (SRA) and can be found under BioProject accession number PRJNA705744. The vcf file for the population genome data can be found in Dryad (https://doi.org/10.25338/B89P86). All data generated or analyzed during this study are included in the published article and its supplementary files.

## ACKOWLEDGEMENTS

We thank Dr. Graham Coop for advice on population genomics analysis and Ernest Lee for advice on bioinformatics analysis. This project was supported by USDA SCRI 2015-51181-24252 and USDA SCRI 2020-67013-30976; USDA APHIS Cooperative Agreement 17-8130-0194-CA to HJB. KML was supported by UC Davis GSR Award.

## DECLARATION OF INTERESTS

The authors declare no competing interests.

## AUTHOR CONTRIBUTION

K.M.L., A.A., Y.L. and J.C.C. conceived the study. A.A. and D.A.W. performed nucleic acid extraction, library preparation and quality control. K.M.L. and W.R.C. performed bioinformatic and population genomics analysis. K.M.L. and J.C.C. wrote the manuscript with the input from all authors. All other authors performed field collections, provided samples for genomic sequencing, and edited the manuscript.

## SUPPLEMENTAL FIGURES

**Supplemental Figure 1:**
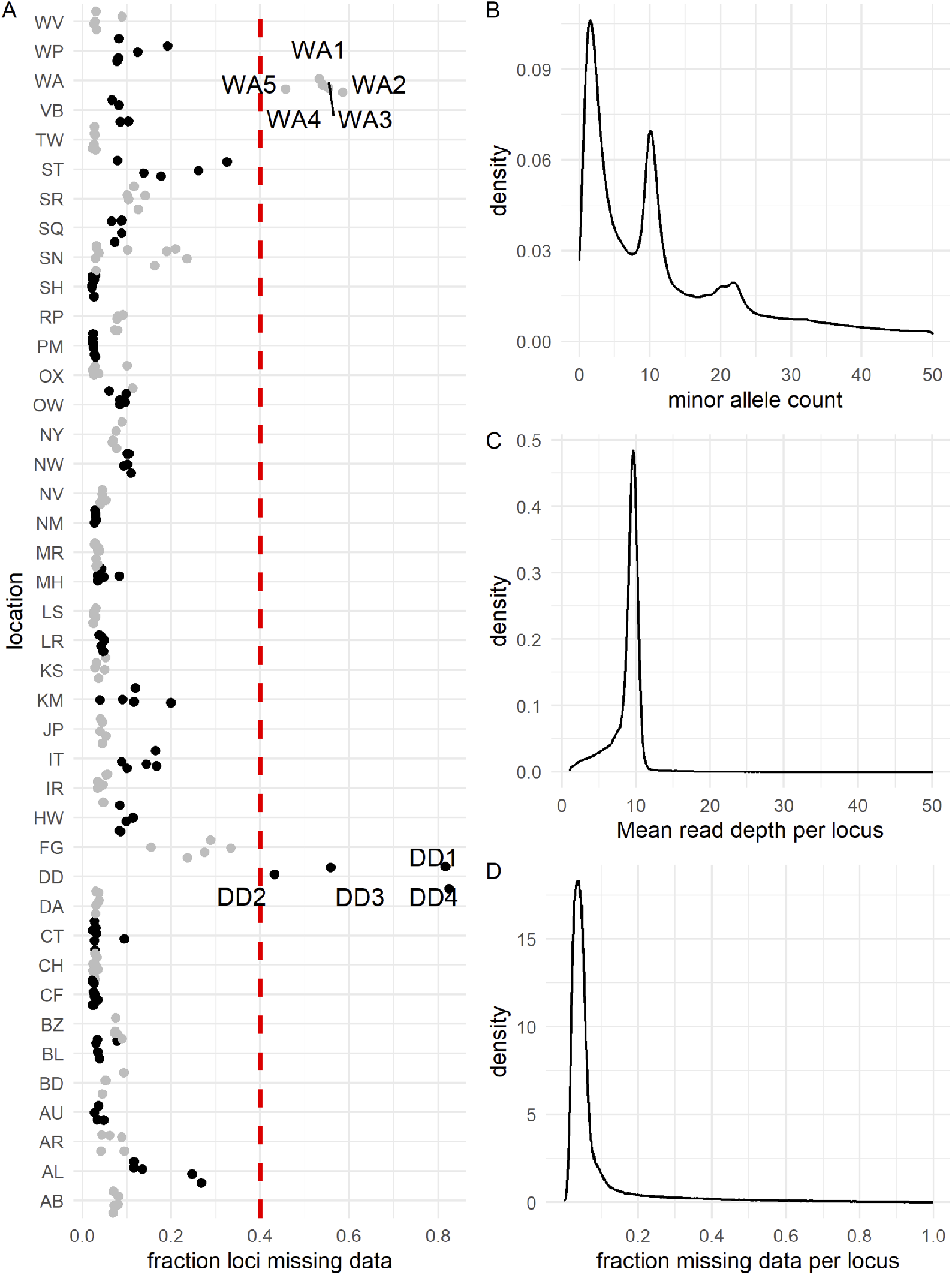
Summary statistics of VCF file prior to filtering for DAPC/Treemix analysis. **(A)** Fraction of loci missing data within each sequenced individual, ordered by sample location. Red line indicates 40% threshold used to exclude samples WA1-5 and DD1-4. **(B)** Minor allele count frequency across all samples. **(C)** Mean read depth per locus frequency across all samples. **(D)** Frequency of the fraction of samples missing data at each locus.

**Supplemental Figure 2:**
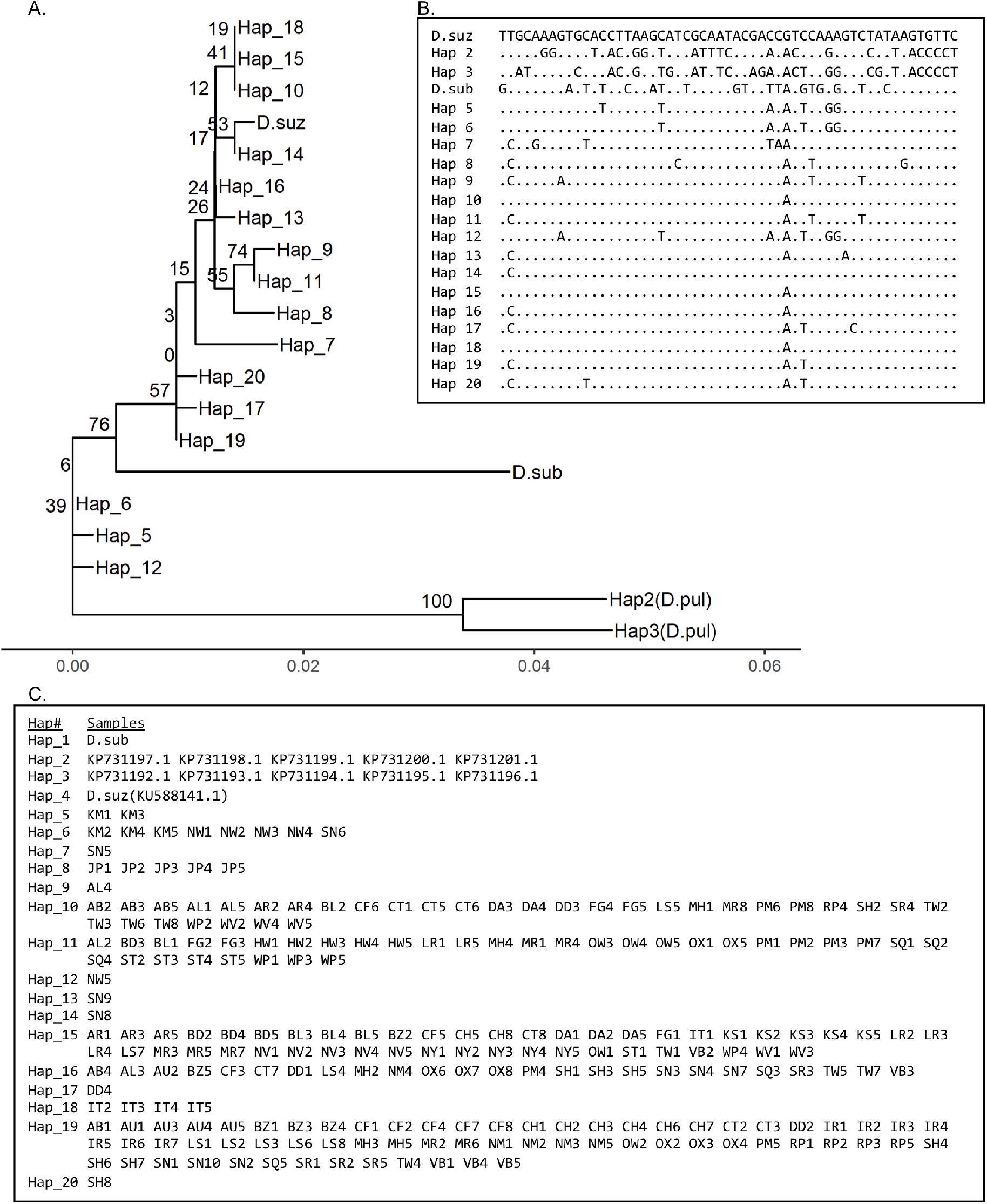
Analysis of COX2 gene between *D. suzukii, D. subpulchrella*, and *D. pulchrella*. **(A)** Maximum likelihood tree of COX2 rooted on haplotype 3, with bootstrap percentages from 500 replicate runs displayed next to branch points. Branch lengths measure number of substitutions per site. 21 nucleotides were analyzed from a total of 595 positions in the alignment. **(B)** Alignment of COX2 gene haplotypes between the *D. suzukii* and *D. subpulchrella* reference mitogenome, ten *D. pulchrella* samples, and the *D. suzukii* samples sequenced for this report. Non-variable sites have been excluded. A period indicates no change from the *D. suzukii* reference. **(C)** List of samples represented by each haplotype.

**Supplemental Figure 3:**
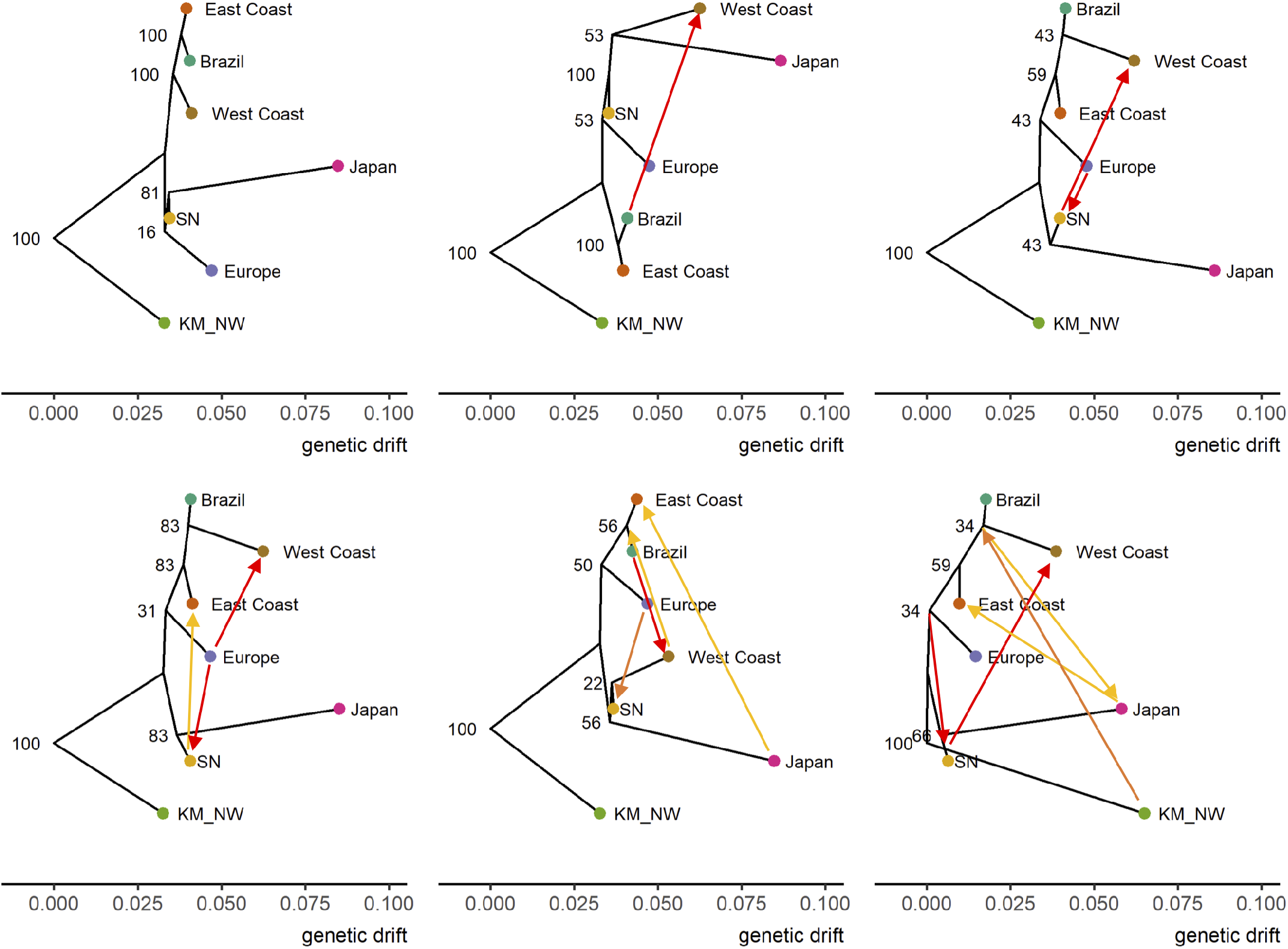
Trees inferred by treemix with 0 to 5 migration edges. Bootstrap replicate values label each branch from 100 bootstrapped runs. Edges are colored by admixture percentage as red (0.35-0.50), orange (0.15-0.34), and yellow (0.01-0.14).

**Supplemental Figure 4:**
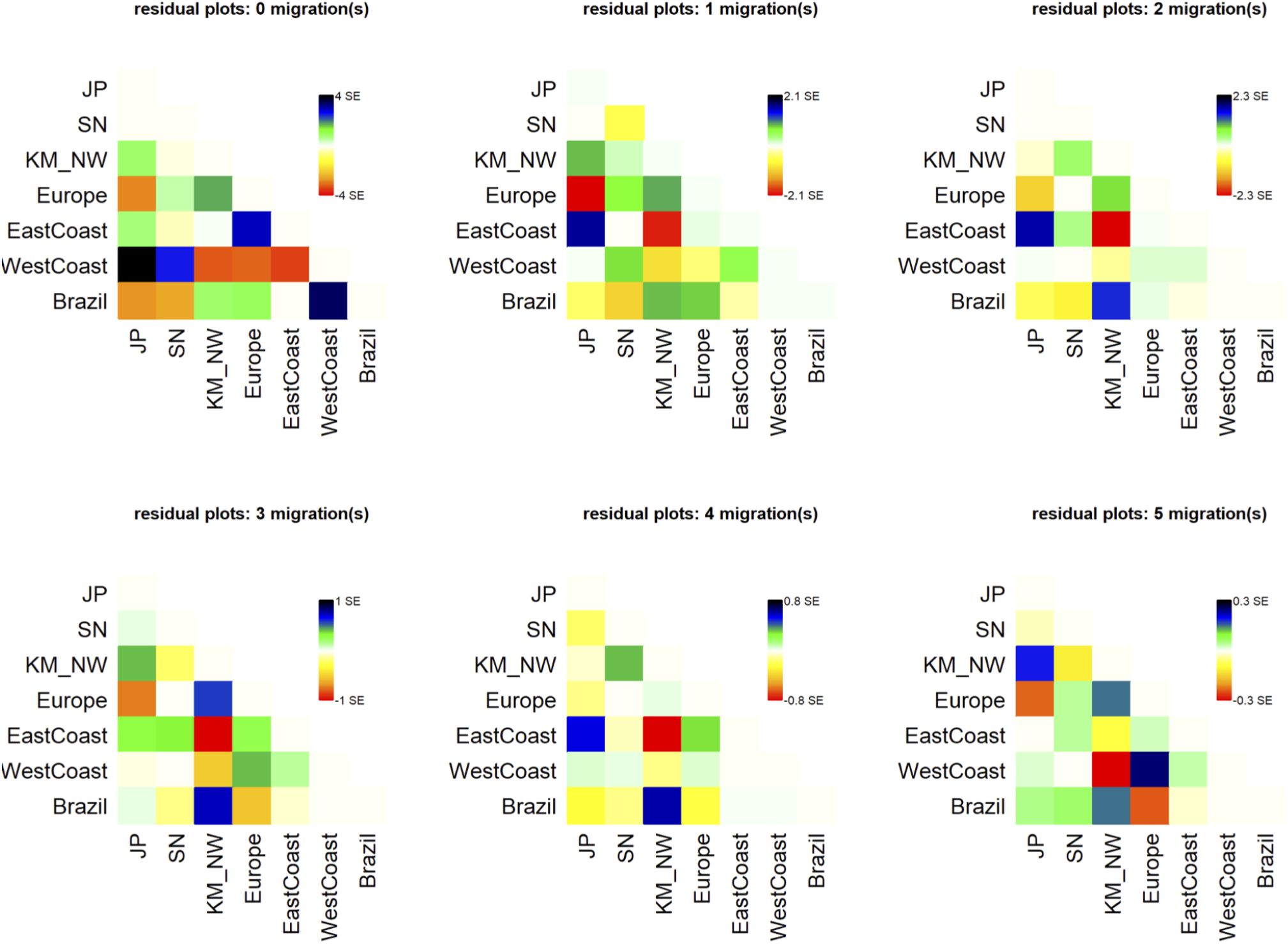
Plots of standard error of residuals between populations based on models generated by Treemix, from 0 to 5 migration edges. High or low values indicate poor fit of the model tree to the data.

## SUPPLEMENTAL TABLES

**Supplemental Table 1:** Locations and time of collection for all *D. suzukii* samples sequenced.

**Supplemental Table 2:** F3 statistics estimated by treemix between all combinations of populations. Significantly negative values for the F3 test (A;B,C) indicate population A experienced gene flow from population B and C.

**Supplemental Table 3**: F4 statistics estimated by treemix between all combinations of populations. Significantly negative values for the F4 test (A,B;C,D) suggest gene flow between A and C or B and D, while positive values indicate gene flow between A and D or B and C.

## Notes

### Competing Interest Statement

The authors have declared no competing interest.

## REFERENCES

Adrion, J. R., A. Kousathanas, M. Pascual, H. J. Burrack, N. M. Haddad et al., 2014 Drosophila suzukii: The genetic footprint of a recent, worldwide invasion. Mol. Biol. Evol. 31: 3148–3163.

Ahmed, H. M. M., F. Heese, and E. A. Wimmer, 2020 Improvement on the genetic engineering of an invasive agricultural pest insect, the cherry vinegar fly, Drosophila suzukii. BMC Genet. 21: 139.

Bergland, A. O., E. L. Behrman, K. R. O’Brien, P. S. Schmidt, and D. A. Petrov, 2014 Genomic evidence of rapid and stable adaptive oscillations over seasonal time scales in Drosophila (D. Bolnick, Ed.). PLOS Genet. 10: e1004775.

Bernt, M., A. Donath, F. Jühling, F. Externbrink, C. Florentz et al., 2013 MITOS: Improved de novo metazoan mitochondrial genome annotation. Mol. Phylogenet. Evol. 69: 313–319.

Bolda, M. P., R. E. Goodhue, and F. G. Zalom, 2010 Spotted Wing Drosophila: potential economic impact of a newly established pest. Agric. Resour. Econ. Update Univ. Calif. Giannini Found. 13: 5–8.

Bolger, A. M., M. Lohse, and B. Usadel, 2014 Trimmomatic: a flexible trimmer for Illumina sequence data. Bioinforma. Oxf. Engl. 30: 2114–2120.

Buchman, A., J. M. Marshall, D. Ostrovski, T. Yang, and O. S. Akbari, 2018 Synthetically engineered Medea gene drive system in the worldwide crop pest Drosophila suzukii. Proc. Natl. Acad. Sci. 115: 4725–4730.

Chang, C. C., C. C. Chow, L. C. Tellier, S. Vattikuti, S. M. Purcell et al., 2015 Second-generation PLINK: rising to the challenge of larger and richer datasets. GigaScience 4: 7.

Chapman, D., B. V. Purse, H. E. Roy, and J. M. Bullock, 2017 Global trade networks determine the distribution of invasive non-native species. Glob. Ecol. Biogeogr. 26: 907–917.

Chen, C.-L., J. Rodiger, V. Chung, R. Viswanatha, S. E. Mohr et al., 2020 SNP-CRISPR: A web tool for SNP-specific genome editing. G3 Genes Genomes Genet. 10: 489–494.

Chiu, J. C., X. Jiang, L. Zhao, C. A. Hamm, J. M. Cridland et al., 2013 Genome of Drosophila suzukii, the spotted wing drosophila. G3 Genes Genomes Genet. 3: 2257–2271.

Cichon, L., D. Garrido, and J. Lago, 2015 Primera deteccion de Drosophila suzukii (Matsumura, 1939) (Diptera: Drosophilidae) en frambuesas del Valle de Rio Negro, Argentina. Libro Resumenes IX Congr. Argent. Entomol. Posadas Misiones 270.

Dalton, D. T., V. M. Walton, P. W. Shearer, D. B. Walsh, J. Caprile et al., 2011 Laboratory survival of Drosophila suzukii under simulated winter conditions of the Pacific Northwest and seasonal field trapping in five primary regions of small and stone fruit production in the United States. Pest Manag. Sci. 67: 1368–1374.

Deprá, M., J. L. Poppe, H. J. Schmitz, D. C. De Toni, and V. L. S. Valente, 2014 The first records of the invasive pest Drosophila suzukii in the South American continent. J. Pest Sci. 87: 379–383.

Dlugosch, K. M., and I. M. Parker, 2008 Founding events in species invasions: genetic variation, adaptive evolution, and the role of multiple introductions. Mol. Ecol. 17: 431–449.

Drury, D. W., A. L. Dapper, D. J. Siniard, G. E. Zentner, and M. J. Wade, 2017 CRISPR/Cas9 gene drives in genetically variable and nonrandomly mating wild populations. Sci. Adv. 3: e1601910.

Durkin, S. M., M. Chakraborty, A. Abrieux, K. M. Lewald, A. Gadau et al., 2021 Behavioral and genomic sensory adaptations underlying the pest activity of Drosophila suzukii. Mol. Biol. Evol. msab048.

Everman, E. R., P. J. Freda, M. Brown, A. J. Schieferecke, G. J. Ragland et al., 2018 Ovary development and cold tolerance of the invasive pest Drosophila suzukii (Matsumura) in the central plains of Kansas, United States. Environ. Entomol. 1–11.

Ferronato, P., A. L. Woch, P. L. Soares, D. Bernardi, M. Botton et al., 2019 A Phylogeographic Approach to the Drosophila suzukii (Diptera: Drosophilidae) Invasion in Brazil. J. Econ. Entomol. 112: 425– 433.

Fraimout, A., V. Debat, S. Fellous, R. A. Hufbauer, J. Foucaud et al., 2017 Deciphering the routes of invasion of Drosophila suzukii by means of ABC random forest. Mol. Biol. Evol. 34: 980–996.

Garnas, J. R., M.-A. Auger-Rozenberg, A. Roques, C. Bertelsmeier, M. J. Wingfield et al., 2016 Complex patterns of global spread in invasive insects: eco-evolutionary and management consequences. Biol. Invasions 18: 935–952.

Garrison, E., and G. Marth, 2012 Haplotype-based variant detection from short-read sequencing. ArXiv12073907 Q-Bio.

Hauser, M., S. Gaimari, and M. Damus, 2009 Drosophila suzukii new to North America. Fly Times 12–15.

Jakobs, R., T. D. Gariepy, and B. J. Sinclair, 2015 Adult plasticity of cold tolerance in a continental-temperate population of Drosophila suzukii. J. Insect Physiol. 79: 1–9.

Johnson, R. N., and P. T. Starks, 2004 A surprising level of genetic diversity in an invasive wasp: Polistes dominulus in the Northeastern United States. Ann. Entomol. Soc. Am. 97: 732–737.

Jombart, T., and I. Ahmed, 2011 adegenet 1.3-1: new tools for the analysis of genome-wide SNP data. Bioinformatics 27: 3070–3071.

Kaneshiro, K. Y., 1983 Minutes, notes, and exhibitions: Drosophila (Sophophora) suzukii (Matsumura). 24: 179.

Kanzawa, T., 1939 Studies on Drosophila suzukii Mats. 49.

Kolbe, J. J., R. E. Glor, L. Rodríguez Schettino, A. C. Lara, A. Larson et al., 2004 Genetic variation increases during biological invasion by a Cuban lizard. Nature 431: 177–181.

Kopelman, N. M., J. Mayzel, M. Jakobsson, N. A. Rosenberg, and I. Mayrose, 2015 CLUMPAK: a program for identifying clustering modes and packaging population structure inferences across K. Mol. Ecol. Resour. 15: 1179–1191.

Korneliussen, T. S., A. Albrechtsen, and R. Nielsen, 2014 ANGSD: Analysis of next generation sequencing data. BMC Bioinformatics 15: 356.

Kumar, S., G. Stecher, M. Li, C. Knyaz, and K. Tamura, 2018 MEGA X: Molecular evolutionary genetics analysis across computing platforms. Mol. Biol. Evol. 35: 1547–1549.

Lee, Y., H. Schmidt, T. C. Collier, W. R. Conner, M. J. Hanemaaijer et al., 2019 Genome-wide divergence among invasive populations of Aedes aegypti in California. BMC Genomics 20: 204.

Li, H., 2013 Aligning sequence reads, clone sequences and assembly contigs with BWA-MEM. ArXiv13033997 Q-Bio.

Li, F., and M. J. Scott, 2016 CRISPR/Cas9-mediated mutagenesis of the white and Sex lethal loci in the invasive pest, Drosophila suzukii. Biochem. Biophys. Res. Commun. 469: 911–916.

Madeira, F., Y. M. Park, J. Lee, N. Buso, T. Gur et al., 2019 The EMBL-EBI search and sequence analysis tools APIs in 2019. Nucleic Acids Res. 47: W636–W641.

Marçais, G., A. L. Delcher, A. M. Phillippy, R. Coston, S. L. Salzberg et al., 2018 MUMmer4: A fast and versatile genome alignment system (A. E. Darling, Ed.). PLOS Comput. Biol. 14: e1005944.

Medina-Muñoz, M. C., X. Lucero, C. Severino, N. Cabrera, D. Olmedo et al., 2015 Drosophila suzukii arrived in Chile. Drosoph. Inf. Serv. 98: 75.

Murphy, K. A., C. A. Tabuloc, K. R. Cervantes, and J. C. Chiu, 2016 Ingestion of genetically modified yeast symbiont reduces fitness of an insect pest via RNA interference. Sci. Rep. 6: 1–13.

Nei, M., and S. Kumar, 2000 Molecular evolution and phylogenetics. Oxford University Press, Oxford?; New York.

Nei, M., T. Maruyama, and R. Chakraborty, 1975 The bottleneck effect and genetic variability on populations. Evolution 29: 1–10.

Noncitrus Fruits and Nuts 2019 Summary, 2020 USDA Natl. Agric. Stat. Serv. 100.

Olazcuaga, L., A. Loiseau, H. Parrinello, M. Paris, A. Fraimout et al., 2020 A whole-genome scan for association with invasion success in the fruit fly Drosophila suzukii using contrasts of allele frequencies corrected for population structure (N. Singh, Ed.). Mol. Biol. Evol. 37: 2369–2385.

Paris, M., R. Boyer, R. Jaenichen, J. Wolf, M. Karageorgi et al., 2020 Near-chromosome level genome assembly of the fruit pest Drosophila suzukii using long-read sequencing. Sci. Rep. 10: 11227.

Peng, F. T., 1937 On some species of Drosophila from China. Annot. Zool. Jpn. 16: 20–27.

Pickrell, J. K., and J. K. Pritchard, 2012 Inference of population splits and mixtures from genome-wide allele frequency data. PLOS Genet. 8: e1002967.

Rašić, G., I. Filipović, A. R. Weeks, and A. A. Hoffmann, 2014 Genome-wide SNPs lead to strong signals of geographic structure and relatedness patterns in the major arbovirus vector, Aedes aegypti. BMC Genomics 15: 275.

Rozas, J., A. Ferrer-Mata, J. C. Sánchez-DelBarrio, S. Guirao-Rico, P. Librado et al., 2017 DnaSP 6: DNA sequence polymorphism analysis of large data sets. Mol. Biol. Evol. 34: 3299–3302.

dos Santos, L. A., M. F. Mendes, A. P. Krüger, M. L. Blauth, M. S. Gottschalk et al., 2017 Global potential distribution of Drosophila suzukii (Diptera, Drosophilidae) (C. Wicker-Thomas, Ed.). PLOS ONE 12: e0174318.

Schmidt, H., T. C. Collier, M. J. Hanemaaijer, P. D. Houston, Y. Lee et al., 2020 Abundance of conserved CRISPR-Cas9 target sites within the highly polymorphic genomes of Anopheles and Aedes mosquitoes. Nat. Commun. 11: 1425.

Schmidt, P. S., L. Matzkin, M. Ippolito, and W. F. Eanes, 2005 Geographic variation in diapause incidence, life-history traits, and climatic adaptation in Drosophila melanogaster. Evolution 59: 1721–1732.

Schöneberg, T., M. T. Lewis, H. J. Burrack, M. Grieshop, R. Isaacs et al., 2021 Cultural Control of Drosophila suzukii in Small Fruit—Current and Pending Tactics in the U.S. Insects 12: 172.

Shearer, P. W., J. D. West, V. M. Walton, P. H. Brown, N. Svetec et al., 2016 Seasonal cues induce phenotypic plasticity of Drosophila suzukii to enhance winter survival. BMC Ecol. 16: 11.

Soria-Carrasco, V., Z. Gompert, A. A. Comeault, T. E. Farkas, T. L. Parchman et al., 2014 Stick insect genomes reveal natural selection’s role in parallel speciation. Science 344: 738–742.

Steck, G. J., W. Dixon, and D. Dean, 2009 Spotted Wing Drosophila, Drosophila suzukii (Matsurmura) (Diptera: Drosophilidae), a fruit pest new to North America: Florida Dept of Agriculture and Consumer Services, 3 p.

Stephens, A. R., M. K. Asplen, W. D. Hutchison, and R. C. Venette, 2015 Cold hardiness of winter-acclimated Drosophila suzukii (Diptera: Drosophilidae) adults. Environ. Entomol. 44: 1619–1626.

Stockton, D. G., A. K. Wallingford, G. Brind’amore, L. Diepenbrock, H. Burrack et al., 2020 Seasonal polyphenism of spotted-wing drosophila is affected by variation in local abiotic conditions within its invaded range, likely influencing survival and regional population dynamics. Ecol. Evol. 10: 7669–7685.

Stockton, D., A. Wallingford, D. Rendon, P. Fanning, C. K. Green et al., 2019 Interactions between biotic and abiotic factors affect survival in overwintering Drosophila suzukii (Diptera: Drosophilidae). Environ. Entomol. 48: 454–464.

Tajima, F., 1989 The effect of change in population size on DNA polymorphism. Genetics 123: 597–601.

Taning, C. N. T., O. Christiaens, N. Berkvens, H. Casteels, M. Maes et al., 2016 Oral RNAi to control Drosophila suzukii: laboratory testing against larval and adult stages. J. Pest Sci. 89: 803–814.

Trask, J. A. S., R. S. Malhi, S. Kanthaswamy, J. Johnson, W. T. Garnica et al., 2011 The effect of SNP discovery method and sample size on estimation of population genetic data for Chinese and Indian rhesus macaques (Macaca mulatta). Primates 52: 129–138.

Tyukmaeva, V. I., T. S. Salminen, M. Kankare, K. E. Knott, and A. Hoikkala, 2011 Adaptation to a seasonally varying environment: a strong latitudinal cline in reproductive diapause combined with high gene flow in Drosophila montana. Ecol. Evol. 1: 160–168.

Walsh, D. B., M. P. Bolda, R. E. Goodhue, A. J. Dreves, J. Lee et al., 2011 Drosophila suzukii (Diptera: Drosophilidae): Invasive pest of ripening soft fruit expanding its geographic range and damage potential. J. Integr. Pest Manag. 2: G1–G7.

Walton, V. M., H. J. Burrack, D. T. Dalton, R. Isaacs, N. Wiman et al., 2016 Past, present and future of Drosophila suzukii: distribution, impact and management in United States berry fruits. Acta Hortic. 87–94.

Willing, E.-M., C. Dreyer, and C. van Oosterhout, 2012 Estimates of genetic differentiation measured by FST do not necessarily require large sample sizes when using many SNP markers. PLOS ONE 7: e42649.

Wu, N., S. Zhang, X. Li, Y. Cao, X. Liu et al., 2019 Fall webworm genomes yield insights into rapid adaptation of invasive species. Nat. Ecol. Evol. 3: 105–115.

